# 3D modeling of thermostable xylanase from Thermotoga naphthophila a member of GH10 family: characterization studies of recombinant xylanase

**DOI:** 10.1101/2024.09.06.611769

**Authors:** Asma Waris, Ali Raza Awan, Muhammad Wasim, Abu Seed Hashmi, Naeem Rashid, Sehrish Firyal, Aisha khalid, Muhammad Tayyab

**Affiliations:** Institute of Biochemistry & Biotechnology, Faculty of Biosciences, University of Veterinary and Animal Sciences, Lahore, Punjab, Pakistan; Riphah International University, Raiwind Road Campus, Lahore, Punjab, Pakistan; School of Biological Sciences, University of The Punjab, Quaid e Azam Campus, Lahore, Punjab, Pakistan; Department of Biology, Garrison University Lahore, Pakistan

**Keywords:** GH10 xylanases, carbohydrate-binding domains, XYL_TN_, *Thermotoga naphthophila*

## Abstract

The current study was planned keeping in view the significance, industrial impact and import of xylanase to Pakistan. In this study, a thermostable recombinant xylanase from *Thermotoga naphthophila* was produced and characterized. The PCR product (1.1 kb) was purified, ligated in the pTZ57R/T and was used for transformation of DH5α cells. The presence of the gene in the recombinant pTZ57R/T was confirmed by restriction analysis. The gene was sub-cloned in pET21a and expression was examined using BL21 CodonPlus (DE3) cells. The recombinant xylanase was expressed as an intracellular soluble protein. SDS-PAGE demonstrated the purified recombinant xylanase as 37 kDa protein. Xylanase showed its optimal activity at 90°C and pH 7. The enzyme was found thermostable and retained 67% activity after an incubation of 1.5h at 90°C in the presence of Mn^2+^. The xylanase activity was enhanced in the presence of Triton X-I00 while the presence of SDS, Tween 20 and Tween 80 showed a declined impact on the activity. Kinetics studies showed the V_max_ and *K*_m_ values of 2313 μmol/mg/min and 3.3 mg/ml respectively. The 3D structure analysis demonstrated the presence of a conserved active site comprised of two glutamate and substrate accommodate sites comprised of + 1, +2 and -1, -2 xylose binding sites in the structure of xylanase. The ability of this thermostable xylanase to work at a wide range of temperatures and pH makes it a suitable candidate for industrial applications.

## Introduction

Xylanases (E.C.3.2.1.8) are glycosidic enzymes responsible for the breakdown of xylan to xylooligosaccharides and xylose. These enzymes are produced by a diverse group of organisms like algae, bacteria, crustaceans, fungi, insects, protozoa and yeast [1]. Xylanases have a broad range of applications in animal feed, biofuel, food, paper & pulp, pharmaceutical industries, textile and waste treatment [2].

Xylanases are distributed into a variety of families considering GH10 and GH11 as major families with a higher number of members [1, 3]. Generally, xylanases are comprised of modular structures having catalytic domains attached with non-catalytic carbohydrate-binding domains (CBMs), via flexible linker regions [4, 5]. Catalytic domain in GH10 family of xylanases comprised of (α/β) _8_ barrel fold. A cleft extending along the entire length of protein in the C-terminal side of ß-strands of the barrel mediates the sugar-binding and is called an active site or substrate-binding cleft [6]. The active site having two conserved catalytic glutamates, one acting as acid/base and the other functioning as nucleophiles are adjacent to C-terminal ends of 4 and 7 ß-strands [7, 8]. Commonly, aromatic amino acids line the inner wall of the cleft and catalytic nucleophiles and acid/base residues are located at the center of the active site [9]. The region of cleft which is responsible for the binding of xylose residues is called subsites. The subsites are given positive (+*n)* and negative (–*n)* numbers depending upon whether they bind the reducing and non-reducing ends of the substrate respectively. The cleavage of xylosidic bonds occurs in the catalytic site situated between -1 and +1 subsites. Amino acids constituting these subsites, have a crucial role in the recognition of substrate and interact with the substrate by hydrogen bonds in the non-reducing region or hydrophobic interaction in the reducing region [10, 11]. Subsites -1, -2 and -3 are called substrate recognition areas, having residues that make abundant interactions with the ligand while +1, +2 and +3 are called product release areas, comprising residues that present few interactions with the ligand. The crystal structure of GH10 xylanases indicated that -1, -2 and +1 subsites in xylanases are highly conserved [12].

The carbohydrate-binding domains (CBMs) do not directly contribute to the catalytic process but play a primary role in carbohydrate recognition and binding. The presence of CBMs allows the enzyme to concentrate on the surface of polysaccharides and improve the catalytic efficiency [13]. According to the carbohydrate-active enzyme database, CBMs are classified into sequenced-based families and 84 families of CBMs are recognized [14, 15]. The CBM that targets the backbone of xylan belongs to the 2, 4, 6, 15, 22, and 36 families [16].

The linker regions are flexible spacers between the domains of modular proteins that exist with variable length and little sequence conservation. They are composed of sequences, highly enriched in hydroxyl amino acids which are extensively O-glycosylated. The glycosylation of hydroxyl amino acids in linker sequences protects from proteolysis. It is also suggested that serine-rich linkers in xylanases may have role parallel to introns in eukaryotes [16].

There is a great demand for xylanases with higher stability in most industries; therefore it is necessary to find a novel thermostable xylanase suitable for the fulfillment of industrial demands. Present study demonstrated the characterization of locally produced recombinant xylanase and highlighted its 3D model with various domains.

## Materials and methods

### Chemicals and reagents

Chemicals and reagents with high purity were used in this study and were procured from Merck (Germany) and Sigma Aldrich (USA).

### Cloning of xylanase gene

Genomic DNA of *Thermotoga naphthophila* was acquired from the German Collection of Microorganism & cell Culture, DSMZ. Nucleotide sequence of xylanase from *Thermotoga naphthophila* RKU-10 (Accession No: CP001839) was utilized for designing XYL-N: TGTTCATATGAAAATATTGC and XYL-C: GCCTCTGGGAAGCCATCA as forward and reverse primers respectively. Primers were designed after the removal of signal peptide comprised of 57 nucleotides and the forward primer had *Nde* I restriction site. The amplified xylanase gene was cloned in pTZ57R/T by TA cloning and the positive clones were screened based on blue/white screening [17]. Alkaline lysis method was utilized for isolation of plasmid DNA and restriction digestion with *Hin*d III and *Nde* I for confirmation of insert in recombinant pTZ57R/T.

### DNA sequencing and phylogenetic analysis

The restriction confirmed positive clone was used for DNA sequencing [18]. The deduced amino acid sequence was utilized for homology and comparative analysis by NCBI BLAST. The multiple sequence alignment was done using ClustalW and the phylogenetic tree was developed using TreeView software. A Phylogenetic tree was constructed using previously reported closely related xylanases by the neighbor-joining method to show the relationship of xylanase from the current study.

### Xylanase expression and purification

Regarding expression analysis, the purified xylanase gene after restriction digestion of recombinant pTZ57R/T was ligated to pET21a previously restricted with same restriction enzymes. *E. coli* BL21 CodonPlus (DE3) cells were utilized as expression host. Overnight grown transformed cells of BL21 CodonPlus (DE3) with pET21a having xylanase gene were diluted to 1% and were induced with 0.5 mM IPTG after attaining an OD of 0.4 followed by the incubation at 37°C for 4.5h under shaking conditions. The cells were harvested and resuspended in the 50 mM phosphate buffer (pH 7). Cells were lysed by sonication and the soluble and insoluble expression was analyzed by SDS-PAGE [19]. The soluble fraction after sonication was utilized for purification by ion exchange and size exclusion chromatography [20]. The purified recombinant xylanase from *Thermotoga naphthophila* was named XYL_TN._

### Xylanase activity assay

The enzyme activity was studied by dinitrosalicylic acid (DNS) method [21]. The activity assay mixture (150 μL) was prepared by mixing 100 μL of 1% beechwood xylan solution prepared in 50 mM phosphate buffer (pH 7) and 50 μL enzyme followed by incubation at 90 °C for 10 minutes. Reaction was terminated with the addition of 1 mL DNS reagent followed by 10 min boiling in water bath. The liberated reducing sugar was calculated by recording the absorbance at 540 nm using a UV-VIS spectrophotometer (Inno-UV2000, InnoTECH, China). One enzyme activity unit was the amount of enzyme for release of one μmol of xylose per minute under assay conditions. Xylose standard curve was utilized for the estimation of xylanase activity units.

## Biochemical characterization

### Effect of temperature on xylanase activity and thermostability

The effect of temperature on the enzyme activity was examined by measuring the XYL_TN_ activity at wide range of temperature from 40 to 100°C using 1% xylan as a substrate. For thermostability studies, recombinant xylanase was incubated at 90°C separately in the absence and presence of Mn^2+^.The samples were withdrawn after every 10 minutes to examine the residual activity [22].

### Effect of pH on xylanase activity

Xylanase activity was observed at various pH ranging from 3 to 10 using 50 mM of each of acetate buffer (3 to 5), phosphate buffer (5 to 7), Tris-HCl buffer (7 to 9) and Glycine/NaOH buffer (9 to 10). The data was utilized for examining the optimal pH for the maximal enzyme activity [23].

### Effect of metal cations and detergents on xylanase activity

The effect of metal cations on enzyme activity was determined to examine the dependency of enzymes on metal cations, the enzyme activity was determined in the presence of 1 mM of various metal cations including Ca^2+^, Mg^2+^, Fe^2^, Zn^2+^, Co^2+^, Cu^2+^, Ni^2+^, and Mn^2+^. XYL_TN_ activity was also studied in the presence of EDTA, non-ionic (Tween-20, Tween-80 and Triton X-100) and ionic (SDS) detergents [24].

### Kinetic studies of xylanase

Regarding kinetic studies, xylanase activity was analyzed under optimal conditions by varying the substrate concentration from 0.1 to 1 mg/mL. The data obtained was utilized for plotting the Lineweaver Burk Plot and was used for the estimation of *K*_*m*_ and V_*max*_ values [25].

### 3D modeling of XYL_TN_

The 3D model of xylanase was developed using the Expasy Swiss-Model tool of the Expasy server (https://swissmodel.expasy.org/) and the 3D structure, substrate binding site, surface model presentation of xylanase and accommodation of substrate in the substrate site in the surface model was built by PyMol.

## Results

### Cloning and phylogenetic analysis of xylanase gene

PCR resulted in the amplification of the 1.1 kb xylanase gene. Single digestion of recombinant pTZ57R/T confirmed the exact size while double digestion with *Nde* I and *Hin*d III resulted in the liberation of a 1.1 kb fragment from the vector. The obtained DNA sequence showed 93% sequence identity with the xylanase gene from *Thermotoga maritima* (CP011107.1), *Thermotoga petrophila* RUK1 (CP000702.1) and *Thermotoga* Sp. RQ2 (CP000969.1), 82% with *Thermotoga neapolitana* (CP000916.1) and *Thermotoga* Sp. RQ7 (CP007633.1).

Based on of amino acids sequence, the XYL_TN_ showed the identity of 93% with xylanase from *Thermotoga meritima*, 92% with *Thermotoga petrophila*, 82% with *Thermotoga* Sp. RQ7, 81% with *Thermotoga neapolitana*, 72% with *Thermotoga caldifentus*, and 70% with *Psudothermotoga hypogea*. The phylogenetic tree confirmed the clustering of xylanase from the current study with various members of the *Thermotoga* Family (Fig 1).

**Fig 1.**
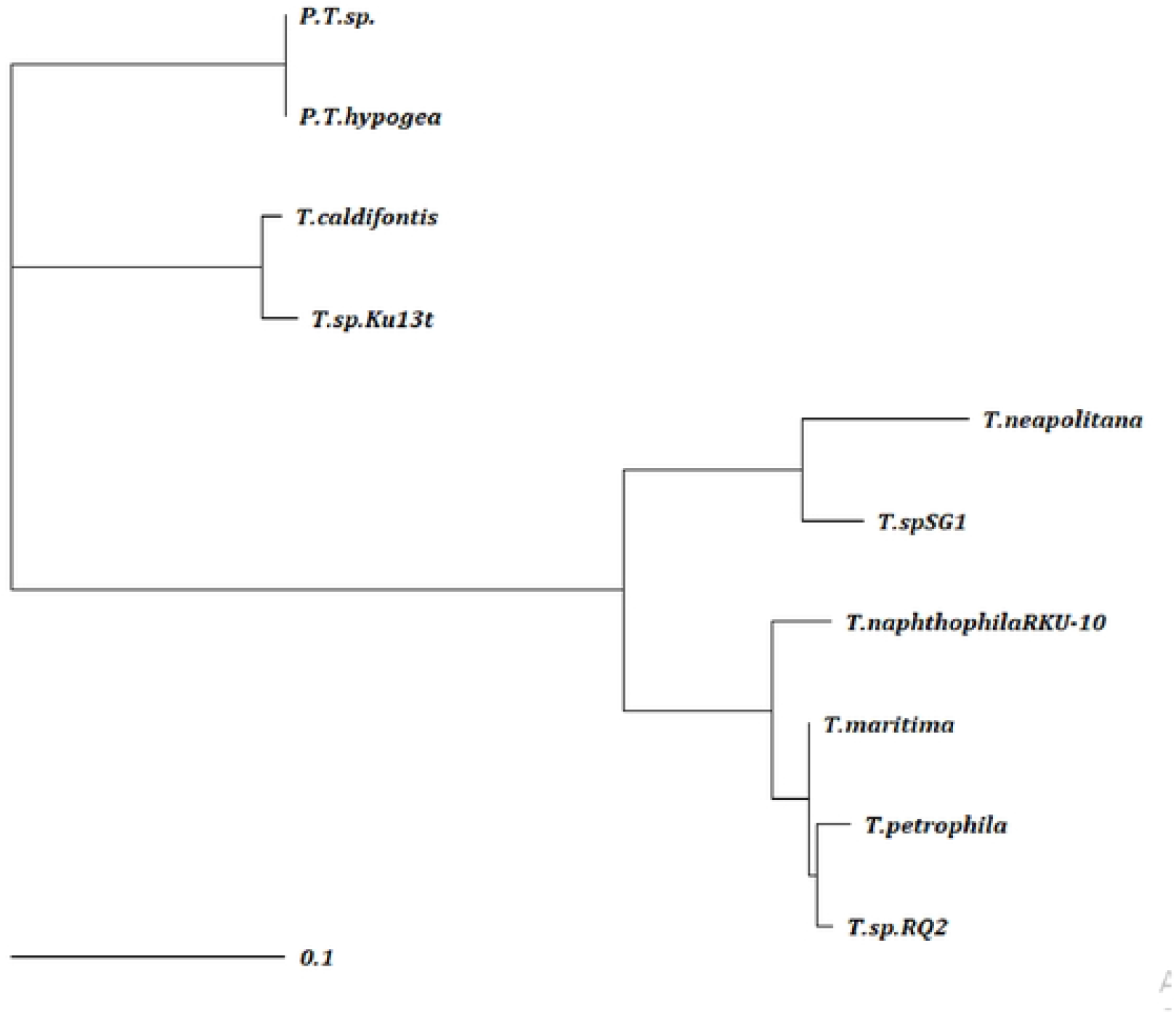
Phylogenetic tree: Tree was constructed based on amino acids sequence of xylanase from *Thermotoga naphthophila* (current study), *P*.*T*.*sp*.: *Pseudo Thermotoga sp*. (MBC7122530), *P*.*T*.*hypogea*: *Pseudo Thermotoga hypogea* (WP081708874), *T*.*caldifontis*: *Thermotoga caldifontis* (WP041077403), T.sp. Ku13t: *Thermotogasp*. Ku13t (KAF2958577), *T*.*neapolitana*: *Thermotoga neapolitana* (WP015919118), *T*.*sp*.*SG1*: *Thermotoga sp*. SG1 (WP101512678), *T*.*meritima*: *Thermotoga meritima* (AAQ01666), *T*.*petrophila*: *Thermotoga petrophila* (WP011943438) and *T*.*sp*.*RQ2*: *Thermotoga sp*. RQ2 (WP012310796) obtained from the NCBI. Name at the end of each branch presents bacterial strain from which xylanases sequences was extracted. Tree was constructed using ClustalW software at the genetic distance 0.1 and was viewed using TreeView software.

### Expression and purification of xylanase

The xylanase was produced as an intracellular soluble protein. The optimal expression of xylanase was observed with the induction of recombinant BL21 CodonPlus (DE3) cells with 0.5 mM IPTG with 5h post-induction incubation at 37°C. SDS-PAGE confirmed the recombinant xylanase as 37 kDa protein (Fig 2).

**Fig 2.**
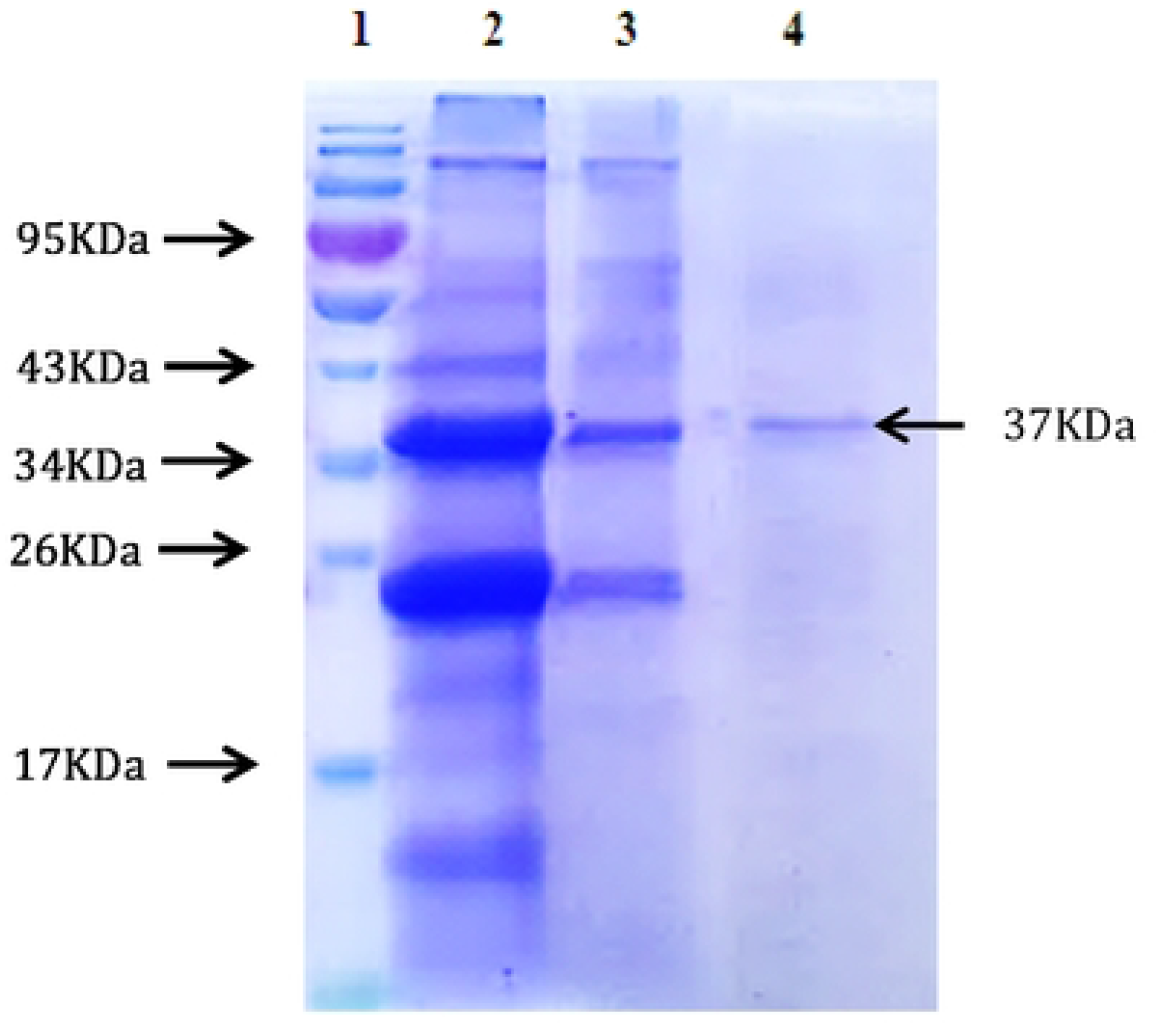
Coomassie Brilliant blue R250 stained SDS-PAGE gel for expression and purification of XYL_TN_. Lane 1: Molecular weight protein ladder (NOVEX, Life Technolgies); Lane 2: Soluble part after lysis of BL21 CodonPlus (DE3) recombinant cells by sonication; Lane 3: Insoluble part after lysis of recombinant cells by sonication. Lane 4: Purified XYL_TN_ after column chromatography.

### Biochemical characterization of XYL_TN_

XYL_TN_ activity was increased with the increase in temperature having maximal activity at 90 °C, whereas a decline in activity was observed beyond 90°C. XYL_TN_ was found stable with enzyme activity > 80 % at 100 °C (Fig 3A). With the increase in pH, XYL_TN_ activity was increased with a maximal activity at pH 7 in 50 mM Tris-HCl buffer (Fig 3B). Thermostability analysis at 90°C showed that the enzyme was stable with 55% or 67% activity after incubation of 1.5h in the absence and presence of Mn^+2^ respectively (Fig 3C). Kinetics studies demonstrated the *K*_*m*_ and V_max_ values of 3.3 mg ml^-1^ and 2313 μmol/mg /min respectively (Fig 3D).

**Fig 3.**
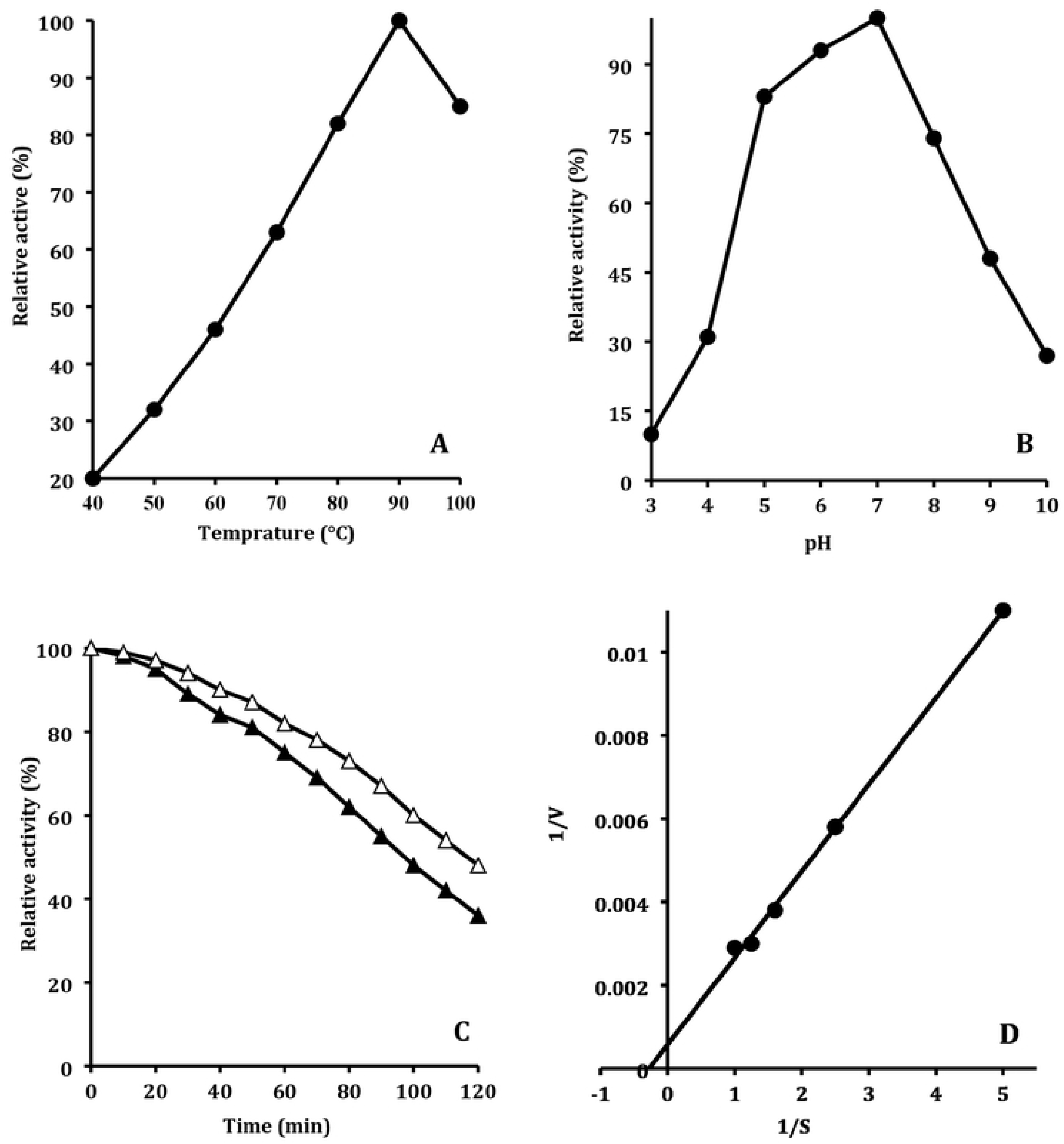
Characterization studies of XYL_TN._ **3A**: Effect of temperature on XYL_TN_ activity. Enzyme activity was performed at various temperatures ranging from 40 to 100 °C using 50 mM Tris-HCl buffer (pH 7). X-axis represents the temperature (°C) and Y-axis shows the relative activity (%). **3B**: Effect of pH on XYL_TN_ activity. Enzyme activity was examined at various pH using 50 mM of each of the Acetate buffer (3-5), Phosphate buffer (5-7), Tris-HCl Buffer (7-9), and Glycine NaOH buffer (9-10) at 90°C. X-axis indicates pH and Y-axis presents XYL_TN_ relative activity (%). **3C**: Thermostability studies of XYL_TN_. Recombinant protein was incubated in the absence and presence of Mn^+2^ separately at 90°C in Tris-HCl buffer (pH 7). The line with close triangles indicates enzyme stability in the absence of Mn^+2^ and open triangles show enzyme stability in the presence of Mn^+2^ ions. X-axis indicates incubation time (min) and Y-axis presents XYL_TN_ relative activity (%). **3D:** Kinetic studies of XYL_TN_. The behavior of xylanase under optimal conditions by varying the substrate concentration from 0.1 to 1 mg/mL. X-axis shows 1/substrate values and Y-axis shows 1/V_0_ values.

Enzyme activity in the presence of 1 mM EDTA indicates that the enzyme is not metal-dependent (Table 1). The enzyme activity was enhanced to 133% or 138% in the presence of 1 mM Co^2+^ or Mn^2+^ respectively. The presence of Fe^2+,^ Cu^2+^, Ca^2+^, Ni^2+^, and Hg^2+^ did not show a significant impact on enzyme activity (Table 1), whereas the presence of Mg^2+^ or Zn^2+^ reduced the xylanase activity to 72% or 73% respectively. The presence of SDS, Tween 80 and Tween 20 reduced the XYL_TN_ activity to 36%, 36% and 50% respectively. However, the presence of 1% Triton X-I00 showed an enhancing impact on XYL_TN_ activity and increased the enzyme activity to 195% (Table 1).

**Table 1.** Effect of Metal cations, EDTA and Detergents on the activity of XYL_TN_.

| **Cations, EDTA and Detergents** | **XYL<sub>TN</sub> Activity (%)** |
| --- | --- |
| **Control** | 100 |
| **EDTA** | 83 |
| **IMPACT OF CATIONS** |  |
| Mn<sup>2+</sup> | 138 |
| Co<sup>2+</sup> | 133 |
| Fe<sup>2+</sup> | 97 |
| Cu<sup>2+</sup> | 96 |
| Ca<sup>2+</sup> | 95 |
| Ni<sup>2+</sup> | 93 |
| Hg<sup>2+</sup> | 92 |
| Zn<sup>2+</sup> | 73 |
| Mg<sup>2+</sup> | 72 |
| **DETERGENTS** |  |
| Triton-X-100 | 195 |
| Tween 20 | 50 |
| Tween 80 | 36 |
| SDS | 36 |

### 3D modeling of XYL_TN_

Amino acids sequence-based comparative analysis of xylanases from the current study with already characterized xylanase from *Thermotoga petrophila* RUK1 showed the presence of conserved catalytic domain active site comprised of two glutamates residues (E^134^ and E^240^); -1 xylose binding site comprises of Lysine (K^52^), Aspargine (N^133^), Glutamine (Q^209^) and Histidine (H^85^); -2 xylose binding site comprises of Glutamic acid (E^48^), Aspargine (N^49^), Lysine (K^52^), Glutamine (Q^92^) and Tryptophan (W^281^); +1 xylose binding site comprises of Tyrosine (Y^177^); and +2 xylose binding site comprises of Serine (S^178^) & Glutamic acid (E^180^) (Fig 4).

**Fig 4:**
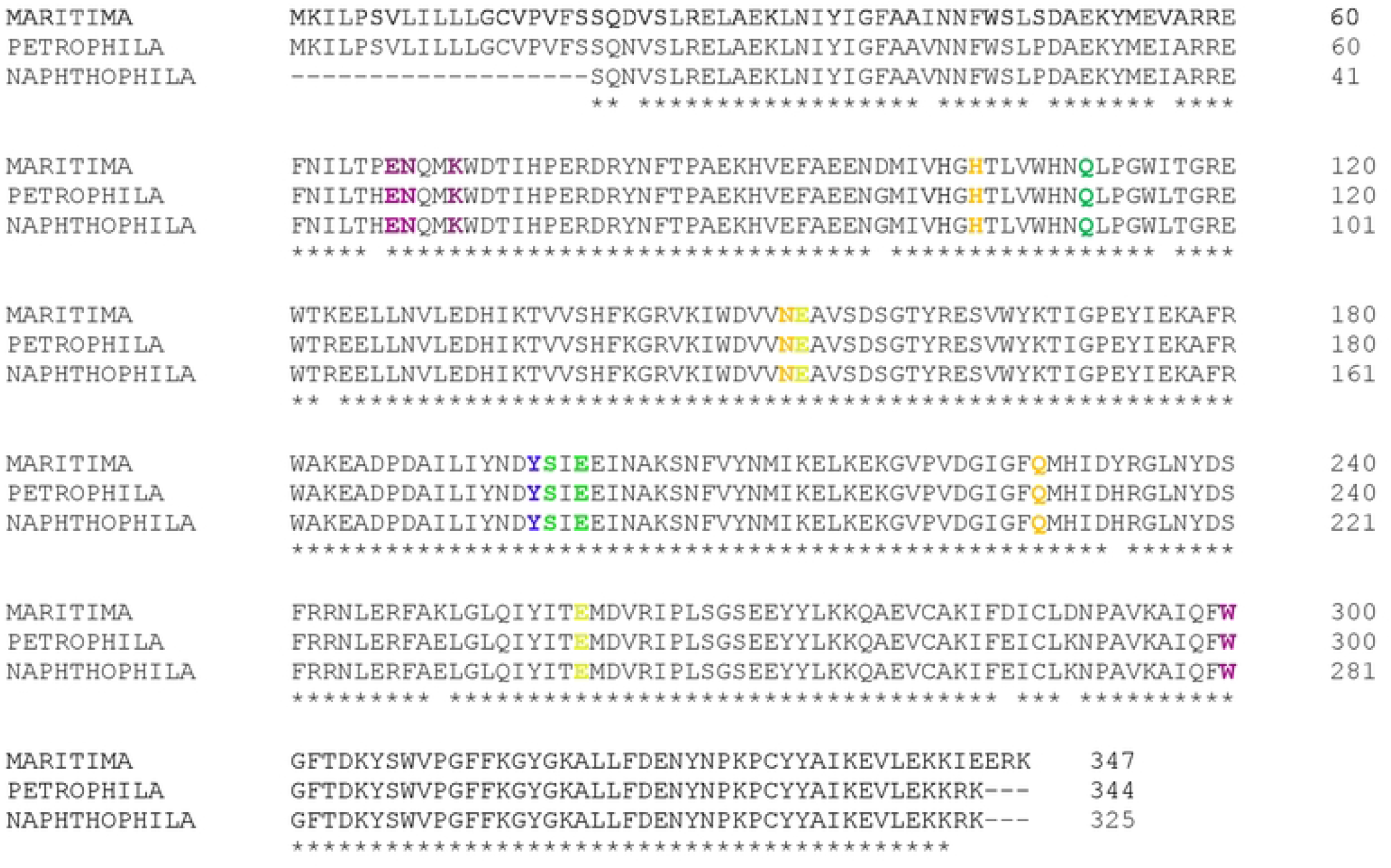
ClustalW was utilized for comparative analysis of the xylanase sequence from *Thermotoga naphthophila* RKU-10 (CP001839.1) with the same enzyme from *Thermotoga petrophila* RUK-1 (CP000702.1) and *Thermotoga maritime* (CP011107.1). The amino acids highlighted in Golden, Pink, Blue, Green and Yellow colors present the -1 xylose binding site (K^52^, N^133^, Q^209 &^ H^85^), -2 xylose binding site (E^48^, N^49^, K^52^, Q^92 &^ W^281^), +1 xylose binding site (Y^177^), +2 xylose binding site (S^178 &^ E^180^), and active site (E^134 &^ E^240^) respectively. K^52^ is part of both -1 and -2 xylose binding sites.

3D model of XYL_TN_ demonstrated the presence of eight parallel β strands forming the barrel structure that is surrounded by the eight α helices structure (Fig 5A). Surface model presentation of XYL_TN_ showed the development of an active site and substrate accommodative groove (Fig 5B) for the accommodation and activity of the enzyme with the help of various conserved domains. The active site and surface accommodative groove without substrate (Fig 5C) and with substrate (Fig 5D) are important for the accommodation of xylooligosaccharide and its breakdown by the enzyme.

**Fig 5.**
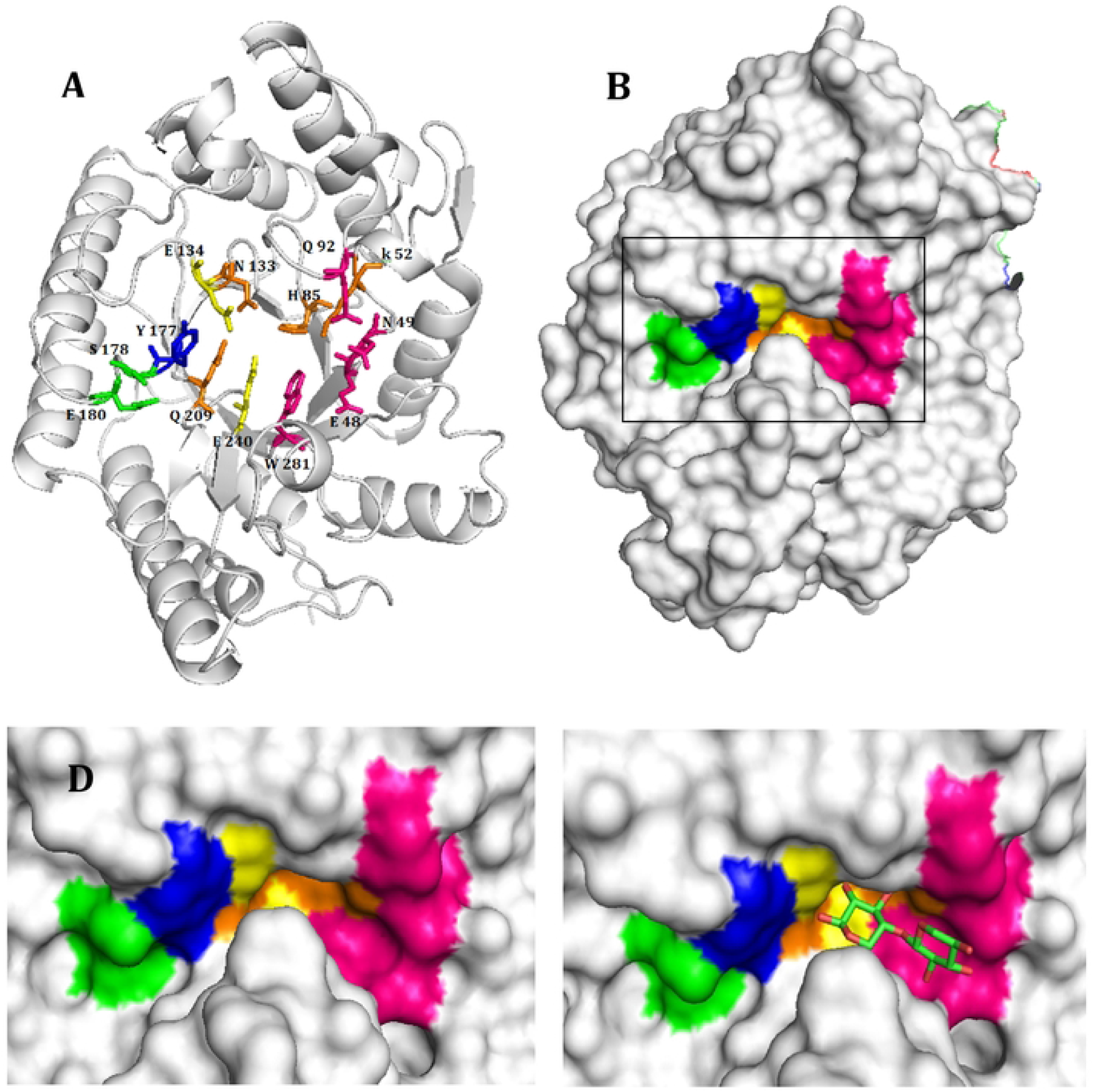
3D structure of XYL_TN_ developed by EXPASY SWISS MODEL. **5A:** Cartoon model for xylanase having highlighted amino acids responsible for binding of substrate and xylanase activity. 5B: Surface Model presentation of xylanase highlighting the pocket of enzyme for interaction with substrate. 5C and 5D are Close views of enzyme pockets for interaction with substrate in the absence or presence of substrate, respectively. The Golden, Pink, Blue, Green and Yellow color presents -1 xylose binding site (K^52^, N^133^, Q^209 &^ H^85^), -2 xylose binding site (E^48^, N^49^, K^52^, Q^92 &^ W^281^), +1 xylose binding site (Y^177^), +2 xylose binding site (S^178 &^ E^180^), and active site (E^134 &^ E^240^) respectively. K^52^ is part of both -1 and -2 xylose binding sites.

## Discussion

Xylanases are depolymerizing enzymes responsible for the hydrolysis of xylan, the second most abundant polysaccharide of plant cell walls to simple absorbable sugars. Xylanases have a wide range of applications including animal feed, baking, bleaching, biofuel, drinks, food, textile, and industries. In the current study, thermostable, recombinant xylanase was characterized from *Thermotoga naphthophila*.

The xylanase showed optimal activity at 90°C which is in agreement with xylanase (XynB) from *Thermotoga maritima* MSB8 [26], whereas xylanases from *Thermotoga thermarum* DSM 5069 [27], *Geobacillus thermoleovorans* [28], and *Thermoanaero bacterium* [29] showed their optimal activity between 70 to 80°C. On the other hand, the xylanase (XynA) from *Thermotoga maritima* MSB8 [26] and xylanases from *Thermotoga thermarum* DSM 5069 [30], *Thermotoga neapolitana* [31] and *Thermotoga sp* FjSS3-B.1 [32] showed their highest activity at temperatures from 92 to 105°C. The optimal activity of XYL_TN_ was recorded at pH 7 which is in agreement with xylanase from *Thermotoga thermarum* DSM 5069 [27]. The xylanase from *Thermotoga thermarum* DSM 5069 [27], xylanase (XynA) and xylanase (XynB) from *Thermotoga maritima* MSB8 [26] showed their maximal activity between pH 5 to 6.2. Whereas xylanases from *Thermoanaero bacterium* [29], and *Geobacillus thermoleovorans* [28] showed their highest activities between pH 7.5 to 8.5, respectively. XYL_TN_ is the metal-independent enzyme that has >80% activity in the presence of 1 mM EDTA. However, the increase in activity due to the presence of Mn^2+^ or Co^2+^ is might be the stabilizing impact of metal ions on the 3D structure of an enzyme. XYL_TN_ demonstrated the V_mex_ of 2313 μmol/mg/min, which is quite high as compared to 374 μmol/mg/min for xylanase (XynA) from *Thermotoga maritima* MSB8 [26], 325.32 μmol/mg/min from *Thermotoga thermarum* DSM 5069 [30], 769 μmol/mg/min from *Thermotoga thermarum* DSM 5069 [30], and 42 μmol/mg/min from *Geobacillus thermoleovorans*, [28] in contrast to this xylanase (XynB) from *Thermotoga maritima* MSB8 [26] and *Thermoanaero bacterium* [29] showed higher level V_max_ of 4600 μmol/mg/min and 3600 μmol/mg/min respectively.

## Conclusion

The current study demonstrated the production and characterization of a recombinant thermostable xylanase from *Thermotoga naphthophila*. The 3D structure and amino acids sequence comparative analysis confirmed the availability and intactness of conserved domains in XYL_TN_ structure that is unique in xylanases of *Thermotoga* family. The ability of this thermostable enzyme to work at a wide range of temperatures and pH makes it a strong candidate to be utilized for industrial applications.

## Author Contributions

**Conceptualization:** Muhammad Tayyab

**Data Curation:** Asma Waris, Aisha Khalid, Muhammad Wasim

**Formal Analysis**: Sehrish Faryal, Naeem Rashid

**Methodology:** Asma Waris

**Supervision:** Muhammad Tayyab

**Writing - original draft:** Asma Waris

**Writing - review & editing:** Muhammad Tayyab, Abu Saeed Hashmi, Ali Raza Awan

